# Functional magnetic resonance imaging of regional impaired cerebrovascular reactivity for migraineurs in the interictal state

**DOI:** 10.1101/859843

**Authors:** Suk-tak Chan, Karleyton C. Evans, Tian-yue Song, Rajiv Gupta, Bruce R. Rosen, Aneesh Singhal, Kenneth K. Kwong

## Abstract

There is still an unmet need of mapping the potential impairment of cerebrovascular reactivity (CVR) in episodic migraineurs in the interictal state. We mapped CVR of 6 episodic migraineurs and 5 headache-free controls (HC) with blood oxygenation level dependent (BOLD) functional magnetic resonance imaging (fMRI) under carbon dioxide (CO_2_) challenge of 30-second epochs with elevated end-tidal partial pressure of CO_2_ (P_ET_CO_2_) by 4-8mmHg. Three migraineurs have migraine without aura (MOA) and the other three have migraine with aura (MA). We found that only MOA subjects showed a reduced or negative BOLD response to CO_2_ at the red nucleus. All 3 MOA subjects were characterized by bilateral posterior communicating artery hypoplasia (bPCAH) identified by MR angiography (MRA). MOA and HC subjects did not show any significant difference in BOLD responses to CO_2_ challenge in cortical and white matter while MA subjects showed poor positive association between BOLD responses and P_ET_CO_2_ in large territories of the cortex and white matter tracts. The combined use of fMRI under CO_2_ challenge and MRA presented a unique approach to investigate the mechanisms of episodic migraine in the interictal state demonstrating for the first time negative CVR at the red nucleus of the midbrain in patients with MOA. CVR maps obtained from both the midbrain and cortical regions provided various signatures to explore the differences between migraineurs and HC and between MOA and MA.

## INTRODUCTION

Since it is rarely possible to image migraineurs in ictal state during migraine attack, imaging the interictal state is a far more realistic proposition. Transcranial Doppler sonography (TCD) under breath hold challenge has been used to evaluate vascular abnormalities in migraine during the interictal period.^1-3^ There had been to our knowledge no functional magnetic resonance imaging (fMRI) study of carbon dioxide (CO_2_) inhalation for migraineurs at the interictal state. There is still a debate on the vascular^4,5^ or neuronal origin of migraine,^6,7^ however, the contribution of vascular disorder to migraine attacks cannot be ignored.^8^ Posterior circulation ischemia has long been hypothesized to be one of the mechanisms for migraine.^9,10^ Visual aura symptoms could be alleviated after the closure of patent foramen ovale,^11^ highlighting a role of blood flow disorder in migraine with aura.

Breath hold challenge has been used as a simple vasoactive stimulus for the assessment of cerebrovascular reactivity (CVR) using TCD in many disorders including carotid artery diseases^12-14^ and brain tumors.^15^ Previously our team used the same technique of combining TCD with breath hold challenge to evaluate vascular abnormalities in migraine during the interictal period.^3^ We found that there was a significant delay in the response of cerebral blood flow velocity (CBFv) to breath hold during 20-second breath holding suggesting a potential impairment of CVR in episodic migraineurs in interictal state. Although TCD offers high temporal resolution to evaluate cerebrovascular responses without the concern of aliasing high frequency hemodynamic signal into the low frequency range, it does not provide regional information. Since fMRI can measure regional hemodynamics with blood oxygen level-dependent (BOLD) signal and a mild hypercapnic challenge with the administration of a low level of CO_2_ can lead to a strong cerebral vascular response with minimal changes in metabolism, heart rate and blood pressure, we propose coupling CO_2_ inhalation with BOLD-fMRI to localize vascular deficits. BOLD-fMRI was used instead of arterial spin labeling (ASL) in MRI perfusion studies due to the low contrast to noise ratio and the low temporal resolution of the ASL technique.^16^ ASL image acquisition at a temporal resolution of 4 seconds will under-sample the brain responses within respiratory cycle of 4-6 seconds.

In the present study, we evaluated if the BOLD-fMRI with mild hypercapnia was able to differentiate migraineurs in interictal state from headache-free controls (HC) in terms of regional CVR. In addition to CVR assessment, the structural integrity of posterior circulation was also studied using magnetic resonance angiography (MRA). We hope that the proposed imaging protocol would provide imaging markers useful for separating migraineurs from non-migraineurs, with additional interest to differentiate migraine with aura (MA) migraine without aura (MOA).

## METHODS

### Subjects

Six right-handed episodic migraineurs and five headache-free controls (9 males, 2 females, aged from 22 to 48 years) were included. Subject demographics are shown in Table 1. Migraine subjects were assessed to have Grade 2 migraine severity using MIGSEV scale.^17^ Three migraineurs had MA while the remaining 3 migraineurs had MOA according to the 2018 IHS 1.1 and 1.2 criteria.^18^ The frequency of migraine attacks was on average 1-10 attacks per month and the duration of the disease was at least one year.

**Table 1.** Demographics of the subjects and the status of posterior communicating arteries (left and right PComm A.) and P1 segment of posterior cerebral arteries (left P1 and right P1). While all the subjects had at least one normal P1 segment of posterior cerebral artery, MOA subjects showed either diminutive or even missing bilateral posterior communicating arteries.

| Subject | Gender | Age | Aura | Frequency of migraine attacks (per month) | Trigger(s) of migraine attacks | Left PComm A. | Right PComm A. | Left P1 | Right P1 |
| --- | --- | --- | --- | --- | --- | --- | --- | --- | --- |
| ***Headache-free Controls (HC)*** |  |  |  |  |  |  |  |  |  |
| S1 | M | 35 | - | - | - | Normal | Normal | Normal | Normal |
| S2 | M | 27 | - | - | - | Normal | Missing | Normal | Normal |
| S3 | M | 32 | - | - | - | Diminutive | Normal | Normal | Missing |
| S4 | M | 22 | - | - | - | Diminutive | Normal | Normal | Normal |
| S5 | M | 32 | - | - | - | Diminutive | Diminutive | Normal | Normal |
| ***Migraineurs without aura (MOA)*** |  |  |  |  |  |  |  |  |  |
| S6 | M | 48 | - | 10 | Alcohol | Diminutive | Diminutive | Normal | Normal |
| S7 | M | 22 | - | 1 | Chocolate and stress | Diminutive | Diminutive | Normal | Normal |
| S8 | M | 38 | - | 5 | Alcohol and stress | Missing | Missing | Normal | Normal |
| ***Migraineurs with aura (MA)*** |  |  |  |  |  |  |  |  |  |
| S9 | M | 28 | Visual | 2 | - | Diminutive | Diminutive | Normal | Normal |
| S10 | F | 26 | Visual | 1 | Alcohol, stress and flashing light | Diminutive | Normal | Normal | Normal |
| S11 | F | 27 | Visual | 1 | Bright and/or flashing light | Normal | Normal | Normal | Normal |
| *Pcomm A., Posterior communicating artery* |  |  |  |  |  |  |  |  |  |
| *P1, P1 segment of posterior cerebral artery* |  |  |  |  |  |  |  |  |  |

MRI scanning was performed on HC and migraineurs in the interictal phase in the Athinoula A. Martinos Center for Biomedical Imaging at the Massachusetts General Hospital of Partners HealthCare. The ‘interictal’ phase is defined as the period outside of migraine when there are no premonitory symptoms 72 hours prior to or 72 hours after a migraine. All the experimental procedures were explained to the subjects, and signed informed consent was obtained prior to participation in the study. All components of this study were performed in compliance with the Declaration of Helsinki and all procedures were approved by Partners Human Research Committee.

### CO_2_ hypercapnic condition

Resting end-tidal partial pressure of CO_2_ (P_ET_CO_2_) was assessed in each subject via calibrated capnograph. Subjects wore nose-clips and breathed through a mouth-piece on an MRI-compatible circuit.^19^ We adjusted the fraction of inspired carbon dioxide to produce steady-state conditions of normocapnia and mild hypercapnia (4-8 mmHg above the subject’s resting P_ET_CO_2_). The CO_2_ challenge paradigm consisted of 2 consecutive phases (normocapnia and mild hypercapnia) repeating 6 times with 3 epochs of 4 mmHg increase and 3 epochs of 8 mmHg increase of P_ET_CO_2_. The normocapnia phase lasted no less than 60 seconds, while the mild hypercapnia phase lasted 30 seconds. The challenge lasted 10 minutes.

When the subject had exogenous CO_2_ challenge, BOLD-fMRI images were acquired. The partial pressure of CO_2_ (PCO_2_) and partial pressure of oxygen (PO_2_) were sampled through the air filter connected with the mouthpiece and the sampled gases were measured by calibrated gas analyzers. The gas analyzers were again calibrated to the barometric pressure of the day of MRI scanning and corrected for vapor pressure. The respiratory flow was measured with respiratory flow head (MTL300L, AdInstruments, Inc., CO, USA) on the breathing circuit via calibrated spirometer (FE141, AdInstruments, Inc., CO, USA). Electrocardiogram (ECG) was measured using Siemens physiological monitoring unit (Siemens Medical, Erlangen, Germany).

### MRI acquisition

MRI brain scanning was performed on a 3-Tesla scanner (Siemens Medical, Erlangen, Germany). The head was immobilized in a head coil with foam pads. Whole brain MRI datasets included: 1) high-resolution T1-weighted 3D-MEMPRAGE images; 2) Time of flight (ToF) sequence; 3) BOLD-fMRI gradient-echo echo-planar-imaging (EPI) images (TR=1450ms, TE=30ms, FOV=220×220mm, matrix=64×64, thickness=5mm, slice gap=1mm) during exogenous CO_2_ challenge. Physiological changes including PCO_2_, PO_2_ and ECG were measured simultaneously with MRI acquisition. All the physiological measurements were synchronized using trigger signals from the MRI scanner. The total duration for each MR session was one hour. BOLD-fMRI images and physiological recordings were stored for offline data analysis.

### Data analysis

The physiological data were analyzed using Matlab R2014a (Mathworks, Inc., Natick, MA, USA). Technical delay of PCO_2_ and PO_2_ was corrected by cross-correlating the time series of PCO_2_ and PO_2_ with the respiratory flow. End inspiration and end expiration were defined on the time series of PO_2_ and PCO_2_. They were verified by the inspiratory and expiratory phases on the time series of respiratory flow. The breath-by-breath P_ET_CO_2_ was extracted at the end expiration of PCO_2_ time series.

MR angiographic images acquired with ToF sequence were interpreted by a neuroradiologist for the patency of posterior circulation. All the BOLD-fMRI data were imported into the software Analysis of Functional NeuroImage (AFNI)^20^ (National Institute of Mental Health, http://afni.nimh.nih.gov) for time-shift correction, motion correction, normalization and detrending. The first 12 volumes in the first 12 time points of each functional dataset, collected before equilibrium magnetization was reached, were discarded. Each functional dataset was corrected for slice timing, motion-corrected and co-registered to the first image of the first functional dataset using three-dimensional volume registration. It was then normalized to its mean intensity value across the time-series. Voxels located within the ventricles and outside the brain defined in the parcellated brain volume using FreeSurfer^21^ (MGH/MIT/HMS Athinoula A. Martinos Center for Biomedial Imaging, Boston, http://surfer.nmr.mgh.harvard.edu) were excluded from the following analyses of functional images. The time-series of each voxel in the normalized functional dataset was detrended with the 5^th^ order of polynomials to remove the low drift frequency. Individual subject brain volumes with time series of percent BOLD signal changes (ΔBOLD) were derived. Linear regression analysis was used to derive CVR by regressing the changes of ΔBOLD on P_ET_CO_2_. The regression coefficient was used to indicate CVR which was defined as the percent BOLD signal changes per mmHg change in P_ET_CO_2_.

The statistical parametric maps for individual subjects were cluster-corrected using a threshold estimated with Monte Carlo simulation algorithm. Individual subject brain volume with CVR magnitude was registered onto each subject’s anatomical scan and transformed to the standardized space of Talairach and Tournoux.^22^ In order to protect against type I error, individual voxel probability threshold of p<0.005 was held to correct the overall significance level to α<0.05. Monte Carlo simulation was used to correct for multiple comparisons.^23^ Based upon a Monte Carlo simulation with 2000 iteration processed with ClustSim program,^24^ it was estimated that a 476mm^3^ contiguous volume would provide the significance level α<0.05, which met the overall corrected threshold of p<0.05.

## RESULTS

CVR maps of all HC, MOA and MA subjects are shown in Figures 1 and 2. Strong positive BOLD responses correlating with the CO_2_ challenge were found throughout the brain in all five HC (Figure 2). BOLD responses to CO_2_ at the cortex and white matter tracts did not show any significant difference between the three MOA subjects and HC (Figure 2). The three MOA subjects showed negative as well as reduced BOLD signal responses to CO_2_ in the red nucleus of the midbrain with one MOA subject (S6) showing bilateral negative BOLD signal change at the red nuclei (Figure 1). This subject also had the highest number of migraine attacks. In all three MOA subjects with negative or reduced BOLD responses at the red nucleus, ToF MR angiography showed bilateral posterior communicating artery hypoplasia (bPCAH), i.e. tiny or no posterior communicating arteries were present (Figure 3). One MA subject (S9) also showed bPCAH, but he did not show reduced or negative BOLD responses to CO_2_ in the red nucleus.

**Figure 1.**
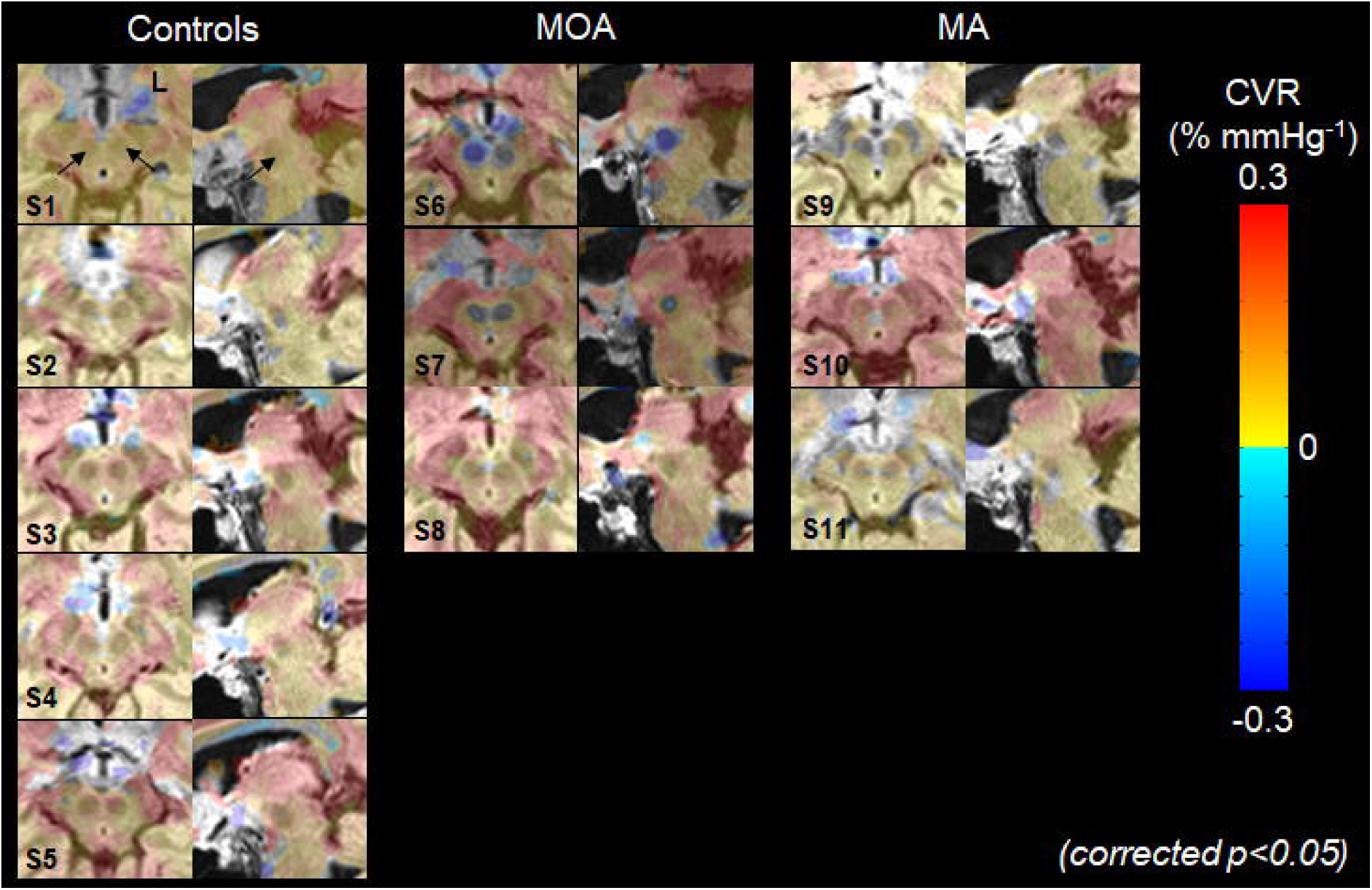
CVR maps of midbrain for headache-free controls (left column), MOA (middle column) and MA subjects (right column). Warm color in the CVR maps represents positive correlation of BOLD signal changes with P_ET_CO_2_ while cold color represents negative correlation of BOLD signal changes with P_ET_CO_2_. Negative and reduced CVR were observed at the red nuclei of MOA subjects.

**Figure 2.**
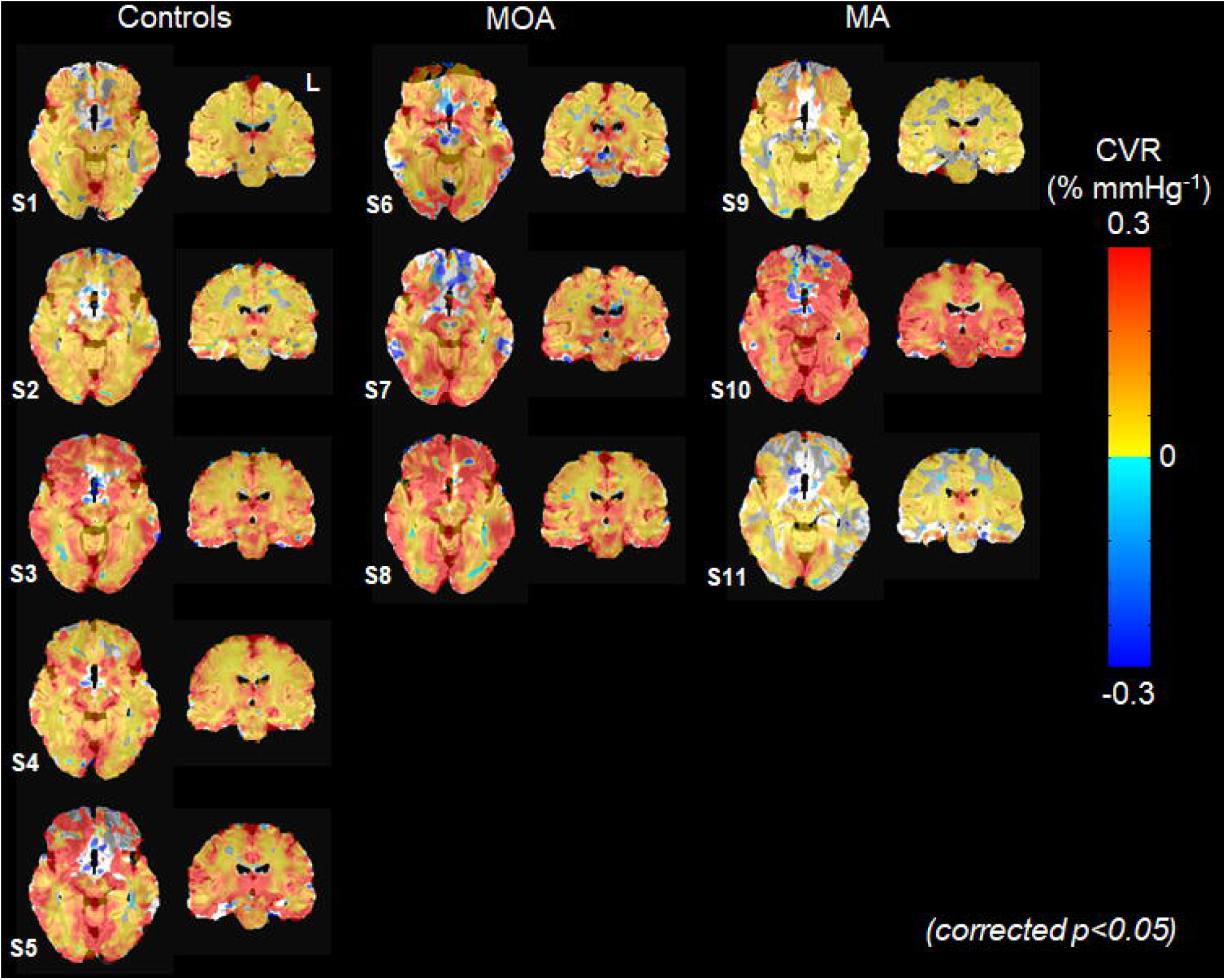
CVR maps of gray and white matter for headache-free controls (left column), MOA (middle column) and MA subjects (right column). Warm color in the CVR maps represents positive correlation of BOLD signal changes with P_ET_CO_2_ while cold color represents negative correlation of BOLD signal changes with P_ET_CO_2_. In two out of three MA subjects, there was a reduced number of brain regions showing significant correlation between BOLD signal changes and P_ET_CO_2_.

**Figure 3.**
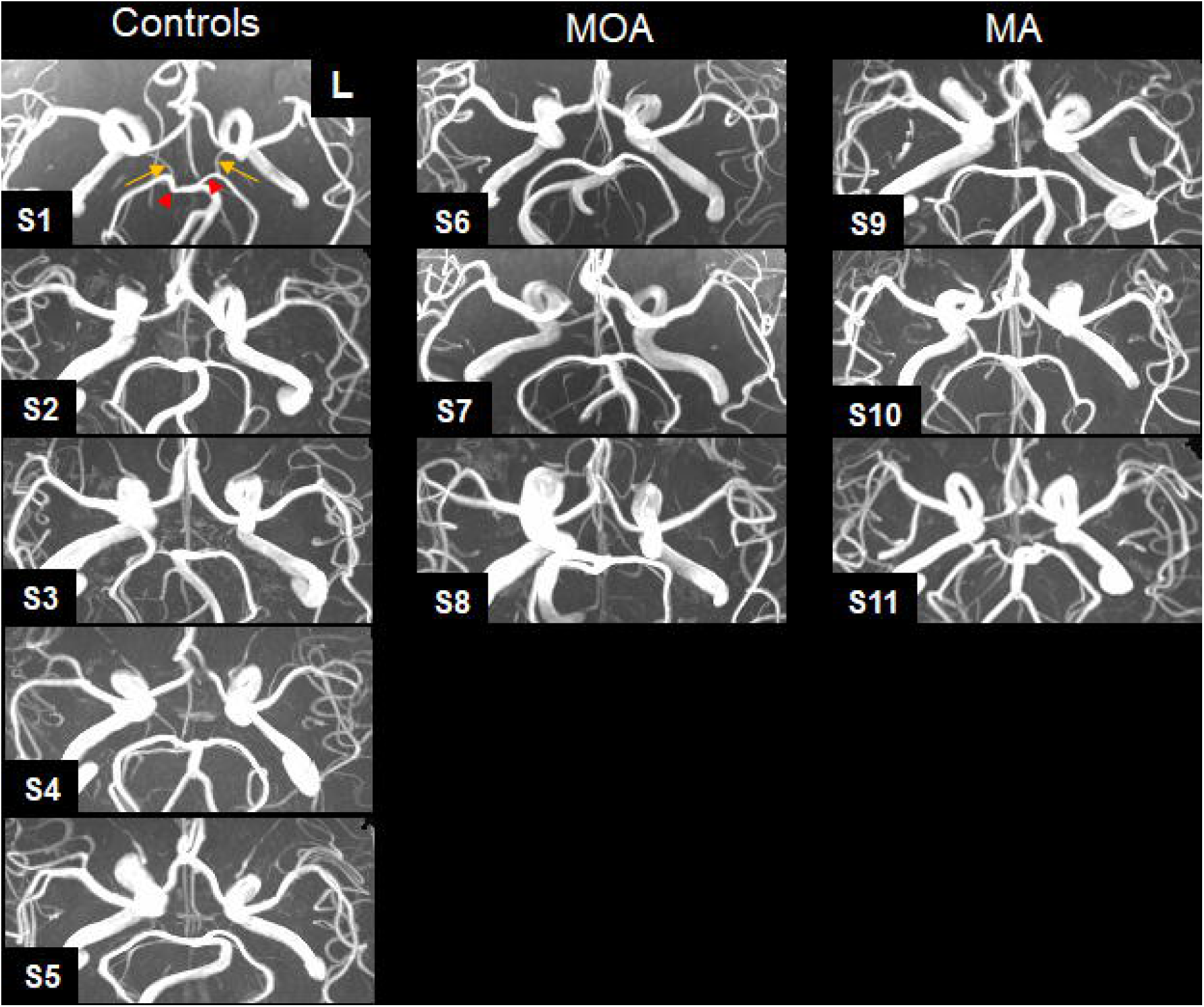
Maximum intensity project (MIP) maps of circle of Willis in headache-free controls (left column), MOA (middle column) and MA subjects (right column). Yellow arrows indicate normal patent posterior communicating arteries and red arrowheads indicate P1 segment of posterior cerebral arteries. In the MOA subject, bilateral posterior communicating artery hypoplasia was observed.

None of the MA or HC subjects showed any reduced or negative BOLD response to CO_2_ at the red nucleus (Figure 4a). All MA subjects, unlike HC or MOA subjects, showed a significantly poor association between BOLD signal changes and measured P_ET_CO_2_ in many gray and white matter regions (Figure 2). Compared to HC or MOA, the total brain volume showing a strong and positive correlation between BOLD signal and P_ET_CO_2_ was much smaller in MA (Figure 4b). In brain areas where there was positive correlation between the BOLD signal and P_ET_CO_2_, the amplitude of CVR was smaller in MA in comparison with MOA and HC, suggesting a more global cerebral impairment of CVR to CO_2_ in MA. None of the migraineurs showed any distinct lesions on FLAIR MRI sequences.

**Figure 4.**
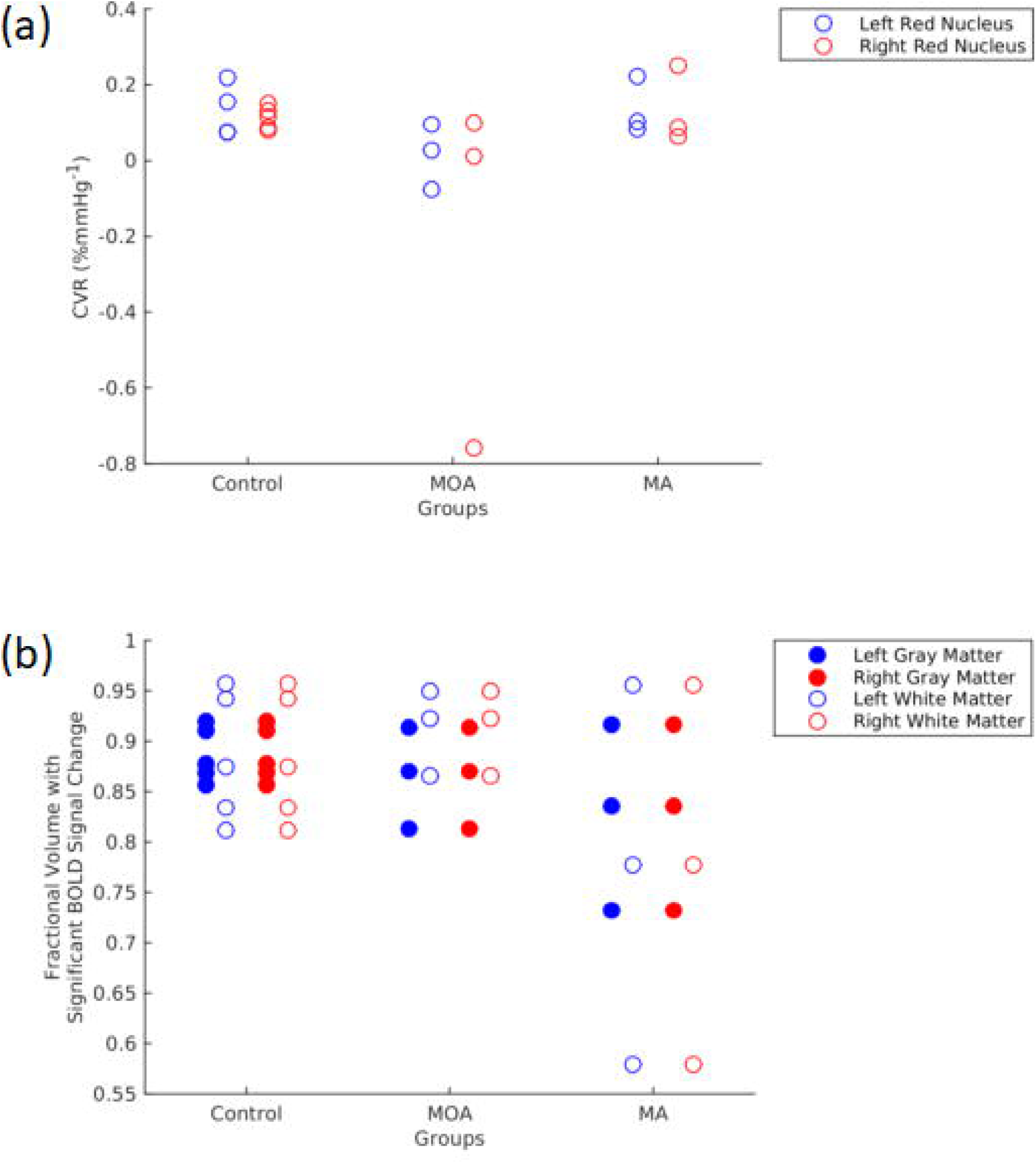
CVR changes in (a) red nuclei and (b) fractional volume of gray and white matter showing significant correlation between BOLD signal changes and P_ET_CO_2_. CVR values at red nuclei in MOA subjects were relatively lower than those in control subjects and MA subjects. Comparing with MOA and control subjects, there was a large variation in the fractional volume of gray and white matter of MA subjects showing significant correlation between BOLD signal changes and P_ET_CO_2_.

## DISCUSSION

Our study shows that significant difference in CVR between HC and migraineurs during the interictal state can be readily identified in individual subjects with low dose CO_2_ challenge. The abnormal CVR observed in gray and white matter for MA but not for MOA subjects or HC, and the reduced or negative BOLD responses to CO_2_ uniquely attributed to MOA subjects at the red nucleus reinforce the hypothesis that MOA and MA are different.

To the best of our knowledge, our finding of reduced or negative CVR at the red nucleus is the only one for any disorders or conditions, and not just for migraine, suggesting abnormal CVR at the red nucleus as a potential marker for migraine. Our HC result was consistent with the findings in another study^25^ of 9 healthy subjects whose BOLD responses at the red nucleus increased under respiratory stress and hypercapnic challenges.

Since all three MOA subjects showed bPCAH, we hypothesize that capillaries distal to the location of hypoplasia supplying the red nuclei were maximally dilated and CO_2_ stress could induce a physiologic “steal” phenomenon diverting blood from the red nucleus to the surrounding normal brain tissue. Such a “steal” phenomenon^26^ under challenges of elevated CO_2_ level and other vasodilators has been reported in stroke,^27^ Moyamoya disease^26^ and traumatic brain injury.^28^ While cited literature presented the general idea of tissue deficits as the hypothesized origin of negative CVR attributed to blood “steal”, our MRI findings offered the opportunity to actually pinpoint the structural condition of bPCAH as an objective candidate to be considered for a mechanism of the blood “steal”.

Besides the presence of bPCAH, presence of increased glutamate may also contribute to the abnormal CVR at the red nucleus of migraineurs. Excessive glutamate presence had been reported at the red nucleus of rats under the condition of neuropathic pain.^29^ MR spectroscopy (MRS) that can correlate elevated glutamate level with abnormal CVR at the red nucleus of migraineurs could be an interesting research target in future.

Since one MA subject also had bPCAH, bPCAH may not be exclusively associated with MOA. With the variable prevalence of bPCAH among MOA and MA subjects,^30^ more studies on CVR to clarify the role of bPCAH as well as red nucleus in MA and MOA are warranted. While reduced or negative CVR at the red nucleus of MOA subjects during the interictal state was shown here for the first time, BOLD signal at the red nucleus^31^ was reported to increase during induced migraine attacks (without any hypercapnic challenge) for both MA and MOA. While it is not surprising that BOLD responses to CO_2_ stress during the interictal state were different from BOLD responses to induced migraine attacks without any involvement of CO_2_ challenge, the common denominator here is that BOLD activities in the red nucleus of migraineurs underwent significant change under a variety of stress conditions. In another MRI study by Kruit et al.,^32^ shortened T2 attributed to iron deposition was observed at the red nucleus of MA and MOA subjects (n=138) in comparison with the T2 of HC (n=75). Collectively these studies showed that the red nucleus was anatomically and functionally different in migraineurs in comparison with controls. It is striking that CVR in the cortex and the white matter did not show significant difference between MOA and HC.

Imaging migraineurs during migraine attacks in the MRI scanner is challenging and an imaging protocol capable of mapping cerebral markers of interest in the interictal state of migraine certainly has its advantage. Our CVR results of individual migraine subjects in the interictal state demonstrated some success in the application of BOLD-fMRI with CO_2_ challenge, while the small sample size of this pilot study was not amenable to pursuing evaluation of any significant group average results. Future studies would be warranted to increase the sample size and to deepen understanding of the preliminary difference observed here between MOA and MA and between migraneurs and headache-free individuals at both the red nucleus and the cortical regions.

## CONCLUSION

The combined use of fMRI under CO_2_ challenge and MRA presented a fresh approach to investigate the mechanisms of episodic migraine in the interictal state demonstrating for the first time negative CVR at the red nucleus of the midbrain of patients with MOA. CVR maps obtained from both the midbrain and cortical regions provided various signatures to explore the differences between migraineurs and HC and between MOA and MA.

## ABBREVIATIONS

ASL: Arterial spin labeling
BOLD: Blood oxygenation level dependent
bPCAH: Bilateral posterior communicating artery hypoplasia
CBFv: Cerebral blood flow velocity
CO_2_: Carbon dioxide
CVR: Cerebrovascular reactivity;
ECG: Electrocardiogram
fMRI: Functional magnetic resonance imaging
HC: Headache-free controls
IHS: International Headache Society
MA: Migraine with aura
MIGSEV: Migraine severity
MOA: Migraine without aura
MRA: Magnetic resonance angiography
PCO_2_: Partial pressure of carbon dioxide
PO_2_: Partial pressure of oxygen
PComm: Posterior communicating artery
P_ET_CO_2_: End-tidal partial pressure of carbon dioxide
TCD: Transcranial Doppler sonography
ToF: Time of flight

## REFERENCES

1. Dora B, Balkan S. Exaggerated interictal cerebrovascular reactivity but normal blood flow velocities in migraine without aura. Cephalalgia. 2002;22(4):288–290.

2. Silvestrini M, Baruffaldi R, Bartolini M, et al. Basilar and middle cerebral artery reactivity in patients with migraine. Headache. 2004;44(1):29–34.

3. Chan ST, Tam Y, Lai CY, et al. Transcranial Doppler study of cerebrovascular reactivity: are migraineurs more sensitive to breath-hold challenge? Brain Res. 2009;1291:53–59.

4. Bigal ME, Kurth T, Hu H, Santanello N, Lipton RB. Migraine and cardiovascular disease: possible mechanisms of interaction. Neurology. 2009;72(21):1864–1871.

5. Kurth T. Associations between migraine and cardiovascular disease. Expert Rev Neurother. 2007;7(9):1097–1104.

6. Goadsby PJ, Charbit AR, Andreou AP, Akerman S, Holland PR. Neurobiology of migraine. Neuroscience. 2009;161(2):327–341.

7. Moskowitz MA. Neurogenic inflammation in the pathophysiology and treatment of migraine. Neurology. 1993;43(6 Suppl 3):S16–20.

8. Asghar MS, Hansen AE, Amin FM, et al. Evidence for a vascular factor in migraine. Ann Neurol. 2011;69(4):635–645.

9. Caplan LR. Migraine and vertebrobasilar ischemia. Neurology. 1991;41(1):55–61.

10. Shin DH, Lim TS, Yong SW, Lee JS, Choi JY, Hong JM. Posterior circulation embolism as a potential mechanism for migraine with aura. Cephalalgia. 2012;32(6):497–499.

11. Khessali H, Mojadidi MK, Gevorgyan R, Levinson R, Tobis J. The effect of patent foramen ovale closure on visual aura without headache or typical aura with migraine headache. JACC Cardiovasc Interv. 2012;5(6):682–687.

12. Ratnatunga C, Adiseshiah M. Increase in middle cerebral artery velocity on breath holding: a simplified test of cerebral perfusion reserve. Eur J Vasc Surg. 1990;4(5):519–523.

13. Vernieri F, Pasqualetti P, Passarelli F, Rossini PM, Silvestrini M. Outcome of carotid artery occlusion is predicted by cerebrovascular reactivity. Stroke. 1999;30(3):593–598.

14. Silvestrini M, Vernieri F, Troisi E, et al. Cerebrovascular reactivity in carotid artery occlusion: possible implications for surgical management of selected groups of patients. Acta Neurol Scand. 1999;99(3):187–191.

15. Pillai JJ, Mikulis DJ. Cerebrovascular reactivity mapping: an evolving standard for clinical functional imaging. AJNR American journal of neuroradiology. 2015;36(1):7–13.

16. Schmithorst VJ, Hernandez-Garcia L, Vannest J, Rajagopal A, Lee G, Holland SK. Optimized simultaneous ASL and BOLD functional imaging of the whole brain. J Magn Reson Imaging. 2014;39(5):1104–1117.

17. El Hasnaoui A, Vray M, Blin P, Nachit-Ouinekh F, Boureau F. Assessment of migraine severity using the MIGSEV scale: relationship to migraine features and quality of life. Cephalalgia. 2004;24(4):262–270.

18. Headache Classification Committee of the International Headache Society (IHS) The International Classification of Headache Disorders, 3rd edition. Cephalalgia. 2018;38(1):1–211.

19. Banzett RB, Garcia RT, Moosavi SH. Simple contrivance “clamps” end-tidal PCO(2) and PO(2) despite rapid changes in ventilation. J Appl Physiol. 2000;88(5):1597–1600.

20. Cox RW. AFNI: software for analysis and visualization of functional magnetic resonance neuroimages. Comput Biomed Res. 1996;29(3):162–173.

21. Fischl B, Sereno MI, Dale AM. Cortical surface-based analysis. II: Inflation, flattening, and a surface-based coordinate system. Neuroimage. 1999;9(2):195–207.

22. Talairach J, Tournoux P. Co-Planar Stereotaxic Atlas of the Human Brain. New York: Thieme Medical; 1988.

23. Gold S, Christian B, Arndt S, et al. Functional MRI statistical software packages: a comparative analysis. Hum Brain Mapp. 1998;6(2):73–84.

24. Ward LL. Simultaneous inference for fMRI data. In: Biophysics Research Institute, Medical College of Wisconsin; 1997.

25. McKay LC, Critchley HD, Murphy K, Frackowiak RS, Corfield DR. Sub-cortical and brainstem sites associated with chemo-stimulated increases in ventilation in humans. NeuroImage. 2010;49(3):2526–2535.

26. Poublanc J, Han JS, Mandell DM, et al. Vascular steal explains early paradoxical blood oxygen level-dependent cerebrovascular response in brain regions with delayed arterial transit times. Cerebrovasc Dis Extra. 2013;3(1):55–64.

27. Dettmers C, Young A, Rommel T, Hartmann A, Weingart O, Baron JC. CO2 reactivity in the ischaemic core, penumbra, and normal tissue 6 hours after acute MCA-occlusion in primates. Acta Neurochir (Wien). 1993;125(1-4):150–155.

28. Long JA, Watts LT, Li W, et al. The effects of perturbed cerebral blood flow and cerebrovascular reactivity on structural MRI and behavioral readouts in mild traumatic brain injury. J Cereb Blood Flow Metab. 2015;35(11):1852–1861.

29. Yu J, Ding CP, Wang J, et al. Red nucleus glutamate facilitates neuropathic allodynia induced by spared nerve injury through non-NMDA and metabotropic glutamate receptors. J Neurosci Res. 2015;93(12):1839–1848.

30. Cucchiara B, Wolf RL, Nagae L, et al. Migraine with aura is associated with an incomplete circle of willis: results of a prospective observational study. PLoS One. 2013;8(7):e71007.

31. Cao Y, Aurora SK, Nagesh V, Patel SC, Welch KM. Functional MRI-BOLD of brainstem structures during visually triggered migraine. Neurology. 2002;59(1):72–78.

32. Kruit MC, Launer LJ, Overbosch J, van Buchem MA, Ferrari MD. Iron accumulation in deep brain nuclei in migraine: a population-based magnetic resonance imaging study. Cephalalgia. 2009;29(3):351–359.

